# North-westward range expansion of the bumblebee *Bombus haematurus* into Central Europe is associated with warmer winters and niche conservatism

**DOI:** 10.1101/2020.02.15.950931

**Authors:** Paolo Biella, Aleksandar Ćetković, Andrej Gogala, Johann Neumayer, Miklós Sárospataki, Peter Šima, Vladimir Smetana

**Affiliations:** University of Milano-Bicocca, Department of Biotechnology and Bioscience, Piazza della Scienza 2, 20126 Milano, Italy; University of Belgrade, Faculty of Biology, Studentski trg 16, 11000 Belgrade, Serbia; Slovenian Museum of Natural History, Prešernova 20, 1001 Ljubljana, Slovenia; Obergrubstraße 18, 5161 Elixhausen, Austria; Szent István University, Department of Zoology and Ecology, Páter K. u. 1., Gödöllő, H-2100, Hungary; Koppert s.r.o., Komárňanská cesta 13, SK-940 53, Nové Zámky, Slovakia; Tekov museum, Sv. Michala 40, SK-934 69 Levice, Slovakia

**Keywords:** climate change, insect conservation, niche conservatism, niche shift, pollinator, range expansion

## Abstract

Range expansions of naturally spreading species are crucial for understanding how species interact with the environment and build their niche. Here, we studied the bumblebee *Bombus haematurus* Kriechbaumer, 1870, a species historically distributed in the eastern Mediterranean area which has very recently started expanding northwards into Central Europe. After updating the global distribution of this species, we investigated if niche shifts took place during this range expansion between colonized and historical areas. In addition, we have explored which climatic factors have favoured the natural range expansion of the species. Our results indicated that *Bombus haematurus* has colonized large territories in 7 European countries outside the historical area in the period from the 1980s to 2018, a natural expansion over an area that equals the 20% of the historical distribution. In addition, this bumblebee performs generalism in flower visitation and habitat use, although a preference for forested areas emerges. The land-use associated with the species in the colonized areas is similar to the historical distribution, indicating that no major niche shifts occurred during the spread. Furthermore, the component of climate change that favoured this colonization is the warming of winter temperatures and similar warming rates took place during both overwintering and queen’s emergence phases. These findings document a case of natural range expansion due to environmental change rather than due to niche shifts, and specifically they indicate that warmer winters are playing a major role in the process of natural colonization of new areas.

## Introduction

The distribution of species’ populations are being influenced by the climatic change and/or by accidental or deliberate introduction events, and thus several taxa are expanding their distribution (Dukes & Mooney 1999; Hickling *et al*. 2006; Chen *et al*. 2011; Biella *et al*. 2017). As a result, species are often colonizing new territories, and the study of these cases have attracted large efforts from ecologists, policy makers, and stakeholders (Bardsley & Edwards-Jones 2007; Essl & Mauerhofer 2018). In most cases, climate change is suggested as one of the major factors that pushes the spread of species in new areas (Wilson *et al*. 2005; Van der Putten *et al*. 2010; Kerr *et al*. 2015), but most of the evidence collected so far are rather circumstantial (Hickling *et al*. 2006; Martinet *et al*. 2015; Barbet-Massin *et al*. 2018). As a consequence, only a minority of researchers have found which components of climate change is favouring these spreading species (Caminade *et al*. 2012; Betzholtz *et al*. 2013). It is therefore particular urgent to investigate which mechanisms are at the root of the colonization of new territories by naturally spreading and by invasive species.

During their spread, species can suddenly start utilizing environmental components that do not entirely resemble the conditions of historical occurrences. This phenomenon takes place if the niche occupied in the colonized area is just a subspace of the niche realized in the historical area, a phenomenon called niche unfilling (Guisan *et al*. 2014; Polidori *et al*. 2018). Instead, if the realized niche in the new areas does not particularly overlap with the native realized niche, then a niche expansion has occurred (Silva *et al*. 2016). Alternatively, when the species maintains its realized niche even in the newly colonized area, it is a case of niche conservatism (Vetaas *et al*. 2018), and its spread is then favoured by a change in the environment or climate in previously unoccupied areas (Pearman *et al*. 2010; Petitpierre *et al*. 2012). All these mechanisms can largely boost the colonization of new areas, but the prevailing mechanism is still disputed. In fact, while the emerging pattern for invading plants seems to be niche conservatism (Petitpierre *et al*. 2012), niche shifts seems to be the main rule for invading insects (Hill *et al*. 2017).

As wild pollinators are subjected to climate change, bumblebees also have been recently changing their distribution. In some cases, human activities are clearly causing invasions into new territories or reinforcing local wild populations (Fraser *et al*. 2012; Aizen *et al*. 2019). In other cases, it seems that bumblebee species are expanding naturally, since some taxa of the subgenus Pyrobombus are expanding their range and locally increasing their relative abundance (Arbetman *et al*. 2017): it is the case with *B. hypnorum* which is quickly colonizing the British Isles and with four other species now increasing their abundance in Vermont (Goulson & Williams 2001; Crowther *et al*. 2019; Richardson *et al*. 2019). Other euro-mediterranean bumblebees have recently colonized areas north of the Arctic Circle, probably favoured by climate warming (Martinet *et al*. 2015), while the opposite route has been taken by the Taiga bumblebee *B. semenoviellus* which is now expanding westwards to central Europe (Šima & Smetana 2012). On the contrary, in response to multiple stressors and environmental factors several other bumblebee species are either shifting their range uphill, have been decreasing their altitudinal range, have negative population trends, and have experienced local extinctions (Cameron *et al*. 2011; Kerr *et al*. 2015; Biella *et al*. 2017; Ornosa *et al*. 2017).

In this study, we have investigated the recent range expansion of the bumblebee *Bombus haematurus* which has recently been found in several Central European countries as result of natural spread northwards from the central Balkanic area (Józan 1995; Neumayer 2004; Sárospataki *et al*. 2005; Šima & Smetana 2009; Jenič *et al*. 2010; Straka *et al*. 2015). We explored two alternative hypotheses on what factor is currently pushing *B. haematurus* northwards. In particular we investigated (a) whether a shift in niche utilization occurred in this species by occupying habitat features that are different from the historical areas, or (b) whether climate warming is favouring the expansion. In addition, we have updated the distribution of the species both in the historical and newly colonized areas and listed the plants this bumblebee is interacting with.

## Material and Methods

### The species

The historical range of *B. haematurus* was provided by a number of authors, but in most cases with little details. The most complete descriptions are given in Reinig (1974) which lists Kopet-Dagh, Elburs, Talesh, Caucasus, Transcaucasia, northern Anatolia, Southern Romania, Bulgaria, and South Yugoslavia, and in (Skhirtladze 1988) there is the addition of the mountains of Crimea, the Stavropol Plateau, the Greater Caucasus, almost all Georgia, Armenia (in the Lesser Caucasus), west and south of Azerbaijan, and northern Iran (and see Results and Discussion of this study). The altitudinal utilization is different according to the latitude, as this species occurs mainly in the lower altitudinal zone (< 400 m a.s.l.) in central Europe, it spreads also into the lower montane zones (up to 1200 m) in the central Balkans, while from southern Bulgaria through Turkey and Iran it can exceptionally reach altitudes as high as 1500-2400 m a.s.l. (Baker 1996; Özbek 1998; Šima *et al*. 2018). Moreover, it has been suggested that this bumblebee is mainly associated with deciduous forests and that it is polylectic (Reinig 1974; Šima *et al*. 2018). A single colony of this bumblebee can produce 80-400 individuals (von Hagen & Aichhorn 2014; Šima *et al*. 2018).

*B. haematurus* belongs to the Pyrobombus subgenus, and *B. hypnorum* is the closest phylogenetic neighbour according to molecular analyses (Cameron *et al* 2007).

The females are easily distinguishable from other co-occurring species by their remarkable colour pattern, especially in Europe (Gokcezade *et al*. 2017). This applies also to males, although it should be noted that some confusion might superficially arise with exceptionally xanthic (yellowish) morphs of *B. pratorum* (Pittioni 1939; Amiet *et al*. 2017).

### The dataset of occurrences and the analysis of distribution

Several sources were used for retrieving the occurrences of this species (see Appendix S1 in Supporting Information for a list). All the duplicated records or those with unclear toponyms or suspected to be unreliable identifications were excluded from the dataset in order avoid unreliable results (see Appendix S1 in Supporting Information for a note on the unreliable cases). This newly created database enumerates 419 unique sites (i.e. records replicated in time from the same site were not counted).

In addition, we have also listed the plant *taxa* that *Bombus haematurus* foraged, by means of field observations or photographs of this bumblebee in the sites used for making the distribution dataset.

The occurrences were grouped in all subsequent analyses as: (1) records lying in the newly colonized area (north of N-Serbia) and (2) records from the historical area (south of N-Serbia). To investigate the extent of the expansion, we calculated the α-hulls around the occurrences of the historical area and around those of the colonized area, that is a standard way for calculating ranges while correcting for the biases of the convex hulls (Burgman & Fox 2003). The α-hulls were calculated with the package *alphahull* in R (Pateiro-Lopez & Rodriguez-Casal 2009); the alpha value was set as the minimum necessary value for drawing a continuous polygon around all occurrences (α = 4) and it was chosen visually by comparing it with the shape generated by minor values. The areas were calculated after excluding the surface of the seas.

### Macro-habitat and land-use data

The data points of occurrence were thinned in order to avoid the effect of oversampled localities and the autocorrelation of spatial and climatic variables as in (Biella *et al*. 2017; Milić *et al*. 2019). To do that, records lying closer than a distance of 3.5 km were removed from the dataset while keeping only one of these records, with the *enmSdm* package for R (R Core Team, 2017; Smith, 2018); the selected threshold was set in order to additively account for the spatial uncertainity of records (e.g. horizonthal error in GPS), the supposed home-range of bumblebees (Knight *et al*. 2005), and in order to avoid overlapping buffers (see below). The resulting dataset listed 330 records and it was used for the following analyses (78.8% of initial records).

To characterize the macro-habitat where the studied species occurs, we extracted the land-use variables from the GlobCover series of land cover maps, that includes yearly layers from 1992 to 2018 (Arino *et al*. 2010; Kirches *et al*. 2017). Firstly, we drew buffers of 1.5 km radius around the occurrences from the thinned dataset. After that, we assigned a chronology to each buffer based on the year(s) of record from all original occurrences from the not-thinned dataset. Then, we calculated the land-use cover of the buffers containing occurrences recorded between 1980 and 2018 as the size of the cover a given land-use type relative to the area of a buffer. Specifically, on one hand, in case of records occurring after the year 1992 (261 single sites), we extracted the land-use cover the year(s) of record of the species in that buffer. On the other hand, in case of records occurring between 1980 and 1991 (n= 32) for which no landuse layer was available, we used the land-use cover mean of the range 1992-1997, a necessary procedure for avoiding loosing several buffers from the historical area. All records before the year 1980 were excluded for the analyses. In case that a given buffer included more years of records, we calculated the mean value between the land-use covers of each year of record of that buffer. In the following analyses, the land use variables were merged into broader categories as indicated in Table 1, but the underrepresented classes were excluded (i.e. occurring in less than 2.5% of the buffers).

**Table 1.** Grouping of GlobCover land-use classes into broader categories as used in this study, and their presence frequency across buffers.

| GlobCover legend | GlobCover codes | Higher grouping | Grouping code | Frequency across buffers (%) |
| --- | --- | --- | --- | --- |
| Cropland, rainfed | 10 | Cropland | CROP | 14.75 |
| Herbaceous cover | 11 | Cropland | CROP | 15.38 |
| Mosaic cropland (>50%) / natural vegetation (tree, shrub, herbaceous cover) (<50%) | 30 | Mosaic | MOSA | 11.62 |
| Mosaic natural vegetation (tree, shrub, herbaceous cover) (>50%) / cropland (<50%) | 40 | Mosaic | MOSA | 5.25 |
| Mosaic tree and shrub (>50%) / herbaceous cover (<50%) | 100 | Mosaic | MOSA | 8.62 |
| Tree cover, broadleaved, deciduous, closed to open (>15%) | 60 | Forest | FORE | 18.75 |
| Tree cover, needleleaved, evergreen, closed to open (>15%) | 70 | Forest | FORE | 6 |
| Tree cover, mixed leaf type (broadleaved and needleleaved) | 90 | Forest | FORE | 5 |
| Grassland | 130 | Grassland | GRSH | 4.88 |
| Urban areas | 190 | Urban | URB | 9.75 |

To test changes in the macro-habitat of occurrence between colonized and historical area for each land use type, we used geographic areas (i.e., colonized vs historical) as predictor variable in interaction with land use categories, with cover percentage in each buffer of a given land-use class as the response variable. We used generalized linear mixed models with buffer identity as random intercept, and Beta distribution that is the appropriate distribution family for variables being proportions that range between 0 and 1 (Cribari-Neto & Zeileis 2010). We used the *glmmTMB* and the *emmeans* packages for R for the regression and the post-hoc test, respectively (Brooks *et al*. 2017; Lenth 2020).

In addition, we also explored which land-use class was preferred by the bumblebee by testing whether the percentages of land-use cover in each buffer were related to specific types of land use. Specifically, the cover percentages in buffers were the response variables and the land-use type was the categorical predictors in generalized linear mixed models with buffer identity nested within area (i.e. historical and colonized) as random intercept and with Beta distribution, that is the appropriate error distribution type for response variables ranging between 0 and 1 and that are proportions (Cribari-Neto & Zeileis 2010). The *glmmTMB* and the *emmeans* packages for R were used for the regression and the post-hoc test, respectively (Brooks *et al*. 2017; Lenth 2020).

### Climatic data

For the localities of recent colonization (i.e. north of Serbia) included in the thinned dataset, temperature data for each year was extracted using the gridded climatic dataset of the CRU TS v4.01 (Harris *et al*. 2014) and we used the layer with mean monthly near-surface temperature. This climatic dataset was used in several studies focused on the distribution of species (e.g. Case and Lawler 2017; Hill *et al*. 2017). Firstly, we divided the year in phases corresponding to (a) the seasons in the Temperate N-hemisphere and considered those ones relevant for bumblebee life cycle (i.e. “winter”: from November to end of February; “spring”: from March to end of May; “summer”: from June to end of August) or (b) overwintering phase (November, December, and January) and emergence phase (February and March, when queens emerge); specifically we calculated the temperature in the recent time frame as a mean between the value in the recording year and the two previous years, and the temperature before the warming maximum as a mean temperature during the years 1977-1979 in these same localities and for each phase. We averaged three years to avoid possible exceptional events occurring in a specific year. We have calculated the difference in temperatures between these two time frames and plotted them against the Euclidean distance of these records from the upper boundary of the historical occurrence area (i.e. “F.G.” = Fruska Gora, 45.1883 N 19.7184 E). The relationship between these variables was tested with linear regressions, with distance from F.G. as the response and the differences in temperature as the predictor. Each phase of the year was tested separately.

## Results

### Distribution

Our dataset indicate that the historical range of *B. haematurus* includes the region from Serbia to northern Anatolia, the Caucasus/Transcaucasia and northern Iran as far east as the Kopet-Dagh mountains (at the border between Iran and Turkmenistan) (Table 2, Fig. 1). The recent expansion, updated until August 2018, includes 188 sites in 7 countries in the upper Balcanic area and Central Europe (Table 2, Fig. 1). In fact, the first reports outside the historical area were found in the year 1987, when the species was recorded in both Hungary and Croatia. In the current range, the following three previously unpublished records highlight that: the westernmost record is now in Italy, near the border with Slovenia (in Cormons, Gorizia province, leg. & det. P. Biella in 2017, new record for the country, >10 workers and 3 ♀); the northernmost record is in Eastern Moravia (in Zlin, Czech Republic, leg. & det. P. Biella in 2017, 1 ♀); in addition, another significant record is in the Upper Danube Valley near Linz (in Haslach, Linz province, Austria, photo of H. Mitterböck in 2018, det. J. Neumayer, 1 ♂).

**Table 2.** Records of *Bombus haematurus* before and after the year 1980 in each country, with details on the first records in the recently colonized countries.

| Country | Year of first colonization | Legit | Det. | Reference | No. Sites | Before 1980 | After 1980 |
| --- | --- | --- | --- | --- | --- | --- | --- |
| Albania | historic |  |  | (Tkalců, 1969) | 1 | 1 | 0 |
| Austria | 1995 | Z. Józán | Z. Józán | (Józán, 1995; Neumayer, 2004) | 52 | 0 | 52 |
| Azerbaijan | historic |  |  | (Skhirtladze, 1988) | 10 | 3 | 7 |
| Bosnia & Herzegovina | historic |  |  | (Reinig, 1974) | 4 | 2 | 2 |
| Bulgaria | historic |  |  | (Reinig, 1974) | 42 | 36 | 6 |
| Croatia | 1987 | G. Mesaroš | A. Četković | A.C. Pers. Coll. | 1 | 0 | 1 |
| Czech Republic | 2013 | J. Straka et al. | J. Straka | (Straka *et al.*, 2015) | 7 | 0 | 0 |
| Georgia | historic |  |  | (Skhirtladze, 1988) | 21 | 21 | 0 |
| Greece | historic |  |  | (Anagnostopoulos, 2005) | 5 | 2 | 3 |
| Hungary | 1987 | Z. Józán | Z. Józán | (Józán, 1995) | 27 | 0 | 27 |
| Iran | historic |  |  | (Özbek, 1998) | 21 | 10 | 11 |
| Italia | 2017 | P. Biella | P. Biella | (Biella & Galimberti 2020) | 3 | 0 | 3 |
| Macedonia | historic |  |  | (Reinig, 1974) | 7 | 7 | 0 |
| Romania | historic |  |  | (Reinig, 1974) | 14 | 12 | 2 |
| Russia | historic |  |  | (Skhirtladze, 1988) | 5 | 5 | 0 |
| Serbia | historic |  |  | (Reinig, 1974) | 66 | 14 | 52 |
| Slovakia | 2003 | V. Smetana | V. Smetana | (Šima & Smetana, 2009) | 83 | 0 | 83 |
| Slovenija | 2007 | J. Grad | A. Gogala | (Jenič *et al.*, 2010) | 15 | 0 | 15 |
| Turkey | historic |  |  | (Reinig, 1974) | 33 | 29 | 4 |
| Ukraine | historic |  |  | (Konovalova, 2010) | 3 | 0 | 3 |

**Figure 1.**
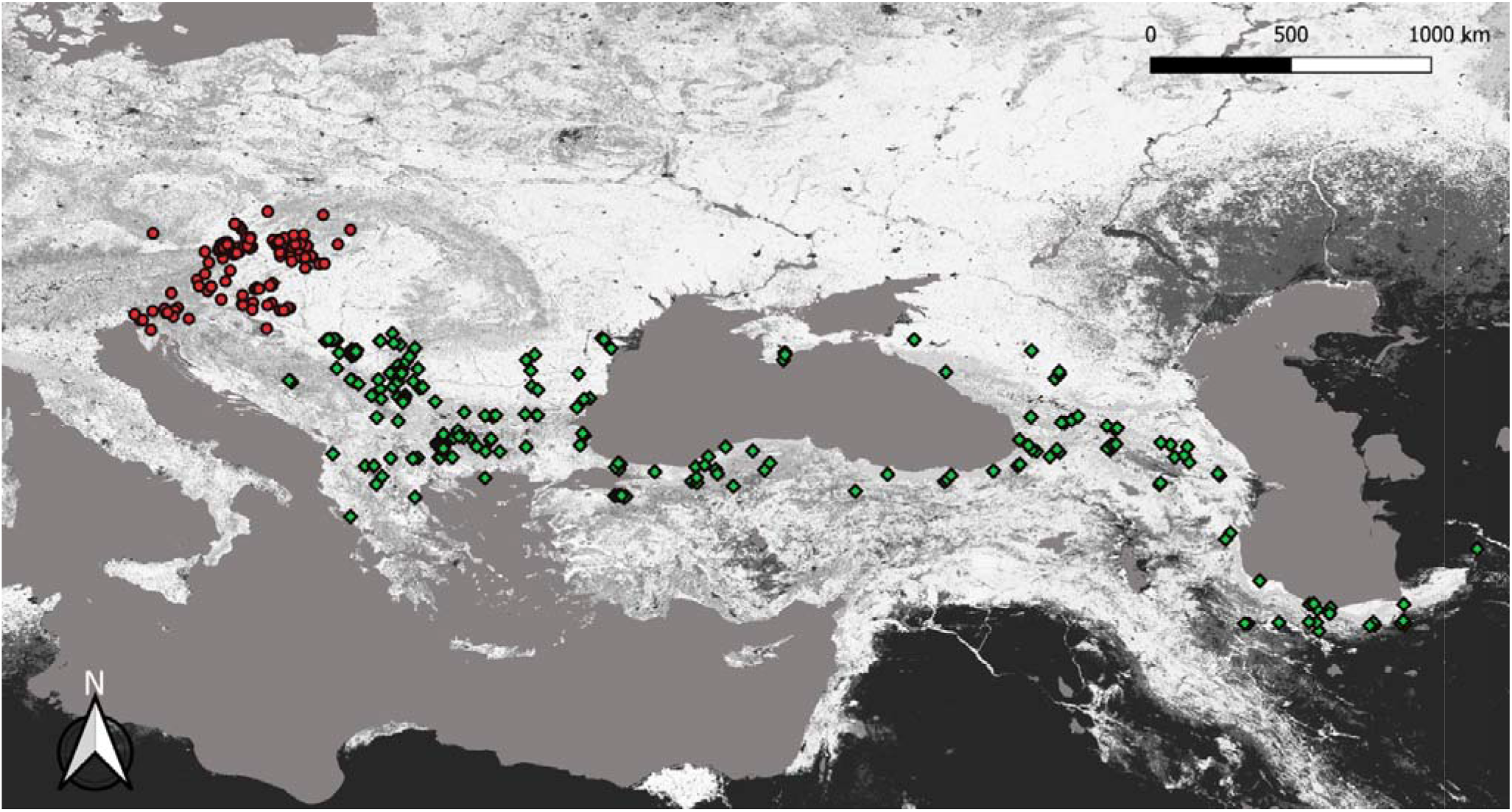
Distribution of the bumblebee *Bombus haematurus*, with red dots indicating the colonized area and the green diamonds indicating the historical area

The α-hull including the current total distribution is 883984 km^2^ (see Figure S1 within Appendix S1 of the Supporting information), and the α-hull of the colonized area corresponds to 21.12% of the historical α-hull (i.e. a colonized α-hull of 154192 km^2^).

### Flower visitation

In our dataset, the bumblebee has been recorded while foraging on a number of plant species with flowers that have open morphology (nectar stored superficially in the corolla) or closed morphology (nectar stored deeply into the corolla) and these plants are herbaceous or woody. Flowers with accessible rewards which were visited by the bumblebee belong to the Boraginaceae family as *Echium vulgare, Borago offinalis, Cerinthe minor, Phacelia tanacetifolia*, and *Lithospermum purpurocaeruleum*, the Liliaceae family as *Fritillaria meleagris*, the Rosaceae family as *Rubus* spp. and *Prunus* spp., the Adoxaceae family as *Viburnum* sp., the Scrophulariaceae family as *Verbascum* sp., and the family Tiliaceae as in *Tilia* (*T. cordata, T*. spp.).

On the other hand, flowers with hidden-nectar morphology are plants of the Lamiaceae family as *Lamium* (L. *album, L. galeobdolon, L. purpureum, L. maculatum), Ballota nigra, Teucrium chamaedrys, Glechoma hederacea, Stachys* (S. *germanica, S. officinalis, S. recta*), *Salvia* (S. *officinalis, S. nemorosa), Lavandula* spp., and *Thymus* spp., the Fabaceae family as *Trifolium pratense, Vicia cracca*, and *Robinia pseudoacacia*, the Boraginaceae family as *Symphytum* (S. *officinalis, S. tuberosum*), the Papaveraceae family as *Corydalis* (C. *cava, C. solida*), the Iridaceae family as *Iris pseudacorus*, the Ericaceae family as *Erica* spp., and the Orobanchaceae family as *Lathraea squamaria*. Additionally, the bumblebee was found while foraging on ornamental plants, such as *Jasminum nudiflorum, Stachys byzantina* and *Rhododendron* hydrids, and other species listed above.

### Habitat use

The buffers around the records of *B. heamoturus* included land-use classes belonging to forest, croplands, mosaic, urban areas, and grasslands (Table 1).

The interaction of land-use type and distribution area variables was a significant predictor in the regression testing changes in the macro-habitat between colonized and historical area (χ^2^=17.318, df=4, p < 0.01). The post-hoc test showed that most of the used land-use categories changed only slightly in their buffers’ cover percentage between colonized areas and the historical areas, with only forests changing significantly (Fig. 2, Table 3).

**Table 3.** Post-hoc test comparing the macro-habitat use between colonized and historical area of *Bombus haematurus*, defined with cover percentages for each land-use type in each occurrence buffer. Statistical significance lower than 0.05 is highlighted in bold and SEM stands for standard error of the mean.

| Contrast: Colonized vs. historical area | Estimate | SEM | P value |
| --- | --- | --- | --- |
| Forest | -0.711 | 0.136 | **<0.001** |
| Cropland | 0.026 | 0.146 | 1.000 |
| Grassland | -0.736 | 0.379 | 0.639 |
| Mosaic | -0.171 | 0.155 | 0.984 |
| Urban | -0.099 | 0.281 | 1.000 |

**Figure 2.**
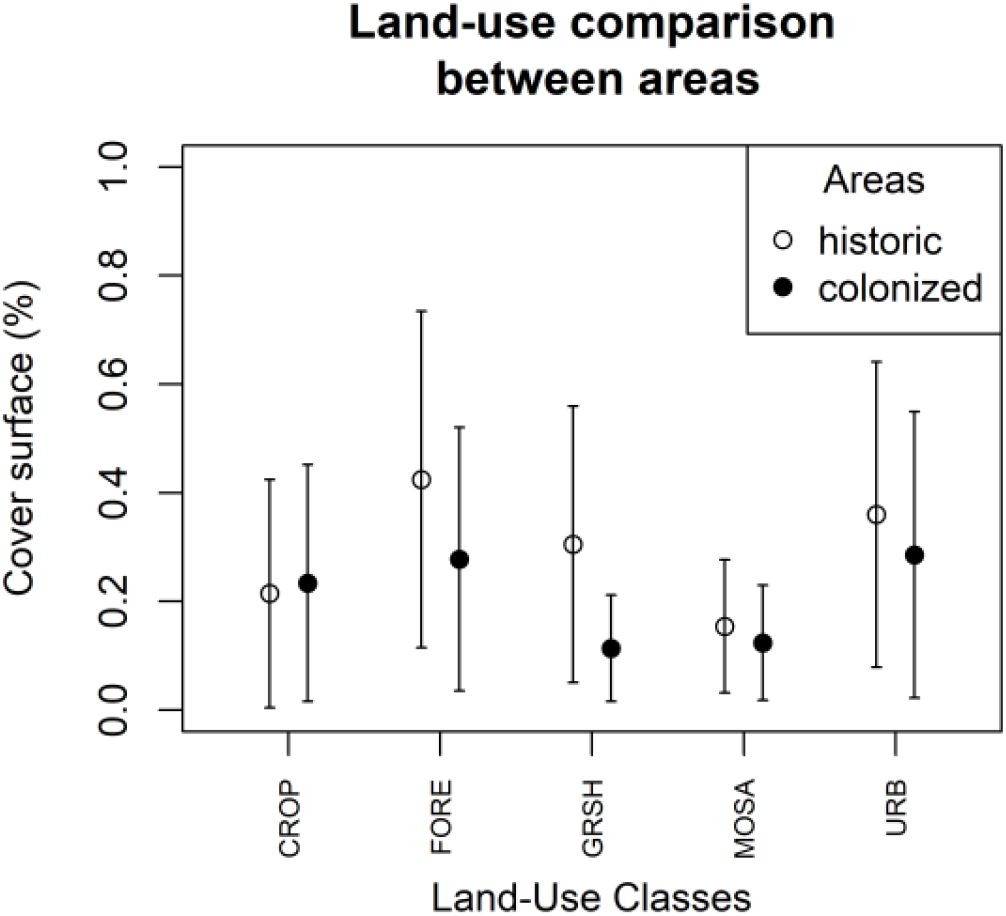
Macro-habitat use by *Bombus haematurus* between colonized and historical areas, expressed as cover percentage in occurrence buffers; see Table 1 for the meaning of the land-use coding in the x-axis.

The land use was a significant predictor in the regression testing the macro-habitat preference (χ^2^=88.47, df=4, p < 0.001). The highest land-use percentages were those of forests, which difference with other categories was significant for most cases except for the urban land-use type, while cropland was significantly higher than mosaic and urban was significantly higher than mosaic (Table 4).

**Table 4.**
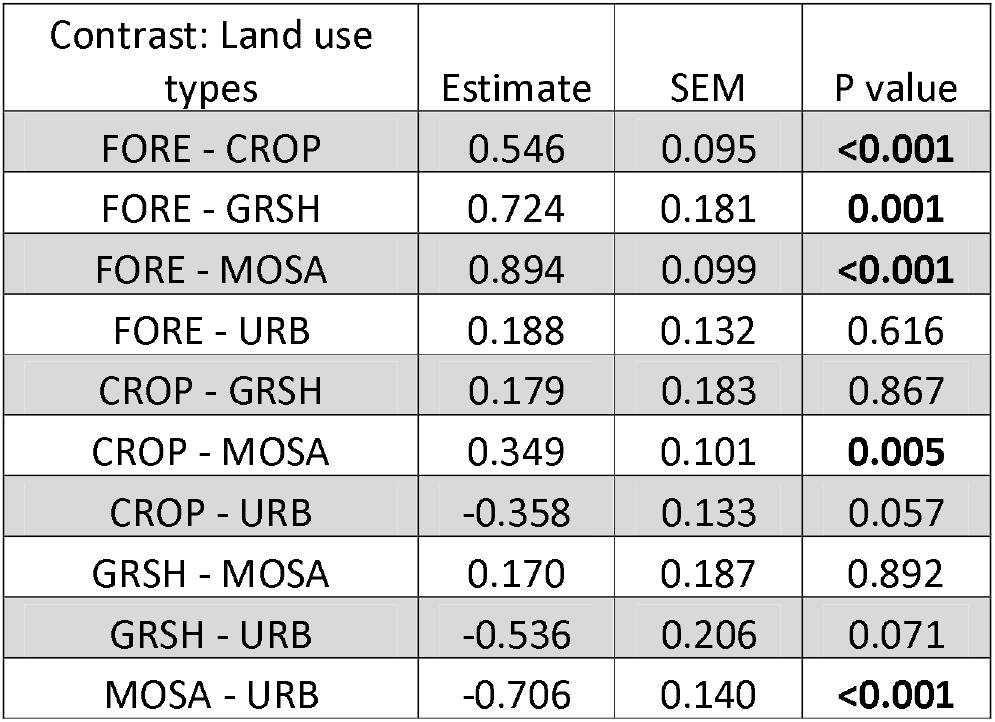
Macro-habitat preference around the occurrences of *Bombus haematurus*, evaluated with a post-hoc test between estimated means of cover percentages for each land-use type; see Table 1 for the meaning of the land-use coding. Statistical significance lower than 0.05 is highlighted in bold and SEM stands for standard error of the mean.

| Contrast: Land use types | Estimate | SEM | P value |
| --- | --- | --- | --- |
| FORE - CROP | 0.546 | 0.095 | **<0.001** |
| FORE - GRSH | 0.724 | 0.181 | **0.001** |
| FORE - MOSA | 0.894 | 0.099 | **<0.001** |
| FORE - URB | 0.188 | 0.132 | 0.616 |
| CROP - GRSH | 0.179 | 0.183 | 0.867 |
| CROP - MOSA | 0.349 | 0.101 | **0.005** |
| CROP - URB | -0.358 | 0.133 | 0.057 |
| GRSH - MOSA | 0.170 | 0.187 | 0.892 |
| GRSH - URB | -0.536 | 0.206 | 0.071 |
| MOSA - URB | -0.706 | 0.140 | **<0.001** |

### Climate change

In the area of colonization, the localities where the bumblebee was recorded were warmer in winter than at the end of the 70s, and this warming was increasing significantly with the distance from northern Serbia (R^2^ = 0.19, p<0.01, Δβ_distance-intercept=0.058_) (Fig. 3). Conversely, the summer phase showed a negative relationship between warming and distance from N-Serbia (R^2^ = 0.027, p<0.05, Δβ_distance-intercept_= −0.044), while during the Spring phase the relationship was not significant (R^2^ = 0.016, P=0.103, Δβdistance-intercept= −0.033) (Fig. 3). Based on the life-cycle of bumblebees, the warming during the overwintering phase was also increasing significantly with the distance from N-Serbia (R^2^ = 0.073, P<0.05, Δβ_distance-intercept_= 0.053), and also warming during the queens’ emergence phase was significantly linear with the distance from N-Serbia (R^2^ = 0.131, P=<0.001, Δβ_distance-intercept_= 0.085) (Fig. 3).

**Figure 3.**
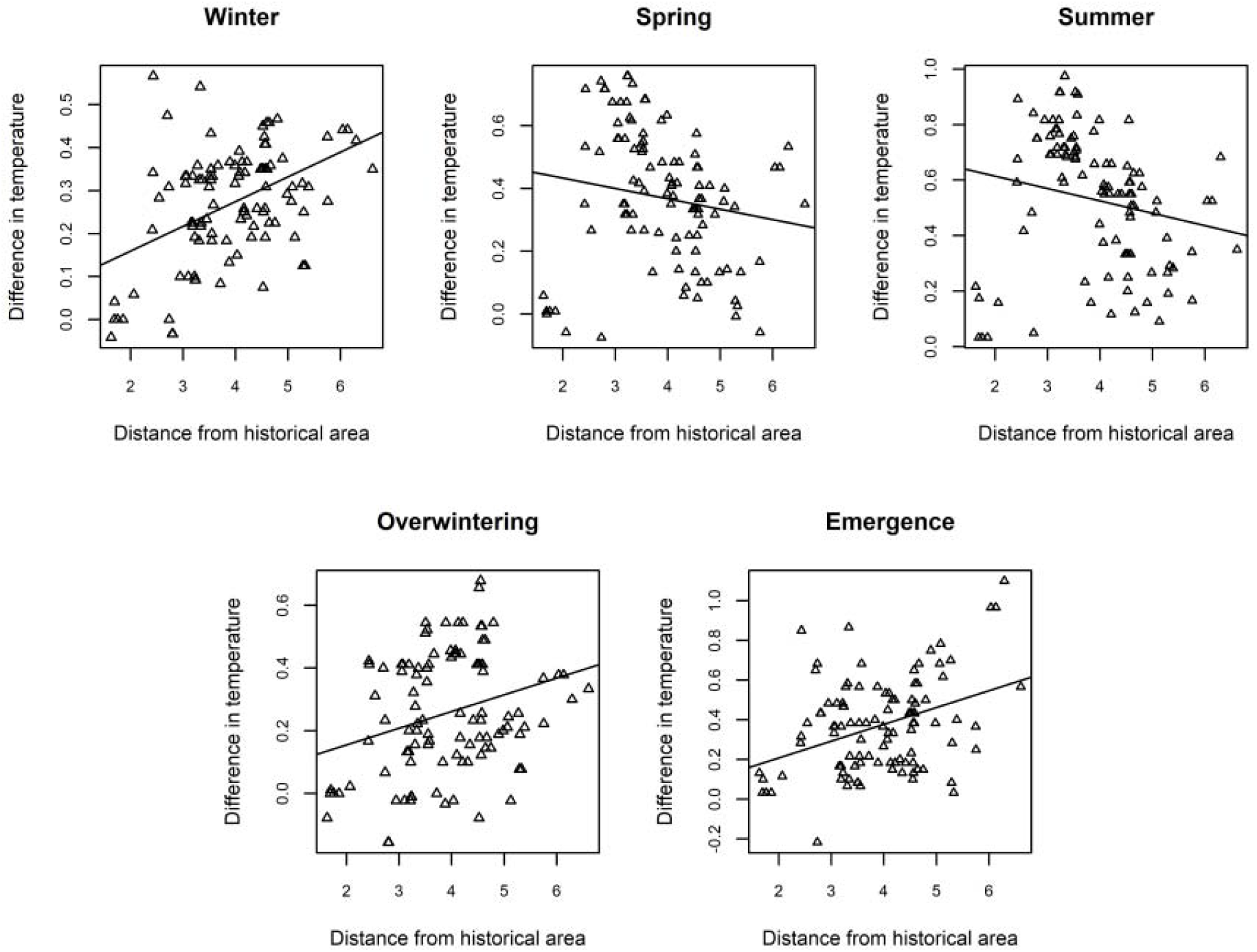
Relation between the temperature change in the colonized area and the distance from the historical area. Differences in temperature between the year of record of *B. haematurus* and the period 1977-1979, separately for winter, summer, and spring and also separately for the overwintering period (Nov. – Jan.) and the queen’s emergence phase (Feb. – Mar.) are represented on the y-axis. On the x-axis, the Euclidean distances between the north boundary of the historical area (Fruska Gora in N-Serbia) and each recorded locality of the recent colonization are shown (it should be noted that the new records lie north of Serbia).

## Discussion

The bumblebee *Bombus haematurus* is now spreading out of its historical range for natural reasons, and during the last few decades, it has largely amplified its range (Šima *et al*. 2018). In this study, we have updated the distribution, investigated the environmental niche of this species and shed light on which components of climate change played a role in favouring ongoing colonization.

Starting in the 1980s, *B. haematurus* has moved northwards and north-westwards from Serbia through Hungary and Croatia and has colonized territories in Central Europe and Italy. Our data showed that the historical area of occurrence extended between N-Serbia and NE-Iran (Fig. 1): a distribution type that should be defined as Balkano-Euxine-Caucaso-Hyrcanian, in accordance with what has previously been suggested for some plant species with a similar distribution (Browicz 1989); considering the real distribution of the studied bumblebee and the regions where it does not occur, this chorological characterization is more appropriate than the more general types suggested in other studies, such as the East-Mediterranean or the Ponto-Mediterranean chorologies (Straka *et al*. 2015; Šima *et al*. 2018). As for the colonized area, in 2017 the northernmost location of the recent area was near the city of Zlin in the Czech Republic, which is significant because it lies in a corridor to north-eastern Europe through the Moravian Gate (i.e. Poland, nearly 70 Km away). In 2018, this species has been recorded in the Upper Danube Valley near Linz in Austria, which leads further north-west into the western part of Central Europe (i.e. Germany, nearly 16.7 Km far). Likewise, a large group of individuals has been recorded in EItaly in 2017, which is the most western occurrence at the edge of an un-colonized vast lowland with hills. In addition, during about 30 years, this bumblebee become quite widespread in the new area since it colonized territories across 7 countries and it was recorded in 188 sites, expanding his distribution over an area equal to about 20% of the historical surface (southern than N-Serbia, that is the area occupied before the 1980s). Given the lack of significant vertical barriers, it is extremely probable that the spread will further continue west, northwest- and northeast-wards in the coming years, which is a prediction further supported by distribution modelling under future climatic scenarios (Rasmont *et al*. 2015).

Rather than niche unfilling or occupying a different niche in the colonized area (Polidori *et al*. 2018), *B. haematurus* is using habitats that are very similar to the historical ones (Fig. 2, Table 3). In addition, we have found that this species inhabits landscapes with forests and with urban surfaces, but it also occurs in mosaics of cropland and natural vegetation, in cropland and in grasslands, both in colonized and historical areas (Fig. 2, Table 3). Although a preference for forested habitat emerges in our data (Table 4), the species is actually occupying several types of habitats; this is in contrast with earlier studies, based on more limited data, which considered this species as a strict hylophilous forest species (Tkalců 1969; Reinig 1971; Šima & Smetana 2009). Our interpretation is that, since the species nests in tree holes (Reinig 1971), queens are likely dependent on mature trees for nesting, but they can use trees outside forests, while foragers disperse into the surrounding landscape for acquiring resources (Šima *et al*. 2018), even in cities (Teppner 2010). As a matter of fact, this bumblebee visits a large number of flowering species with both simple and complex flower shapes (see results for a list), which hints at a generalist strategy (Kaluza *et al*. 2017; Biella *et al*. 2019a, 2019b; Kratschmer *et al*. 2019). The fact that this species is performing a generalist strategy in foraging and that occurs in heterogeneous macrohabitats likely supported this natural spread (Moreyra & Lozada 2020), as this bee is actually exploiting multiple facets of the new environment.

In the extended period prior to the 1980s, *B. haematurus* had a relatively stable north-western range limit in northern Serbia, but there are no records from south-eastern Hungary before the 1980s despite the fact that entomologists were active in this area during that period. Since the colonization of new areas by *B. haematurus* started during the 1980s, we expected climate change to play a major role in this spread. Indeed, the period between the 1980s and the 2010s registered the highest atmospheric temperatures of the last 800 years in the Northern Hemisphere, with a warming rate of 0.2 °C/decade occurring worldwide in the period from 1984 to 1998 (IPCC, Intergovernmental Panel on Climate 2014). Several species responded by shifting their distributive area in that period, such as the alpine bumblebee *Bombus alpinus* that raised its lower altitudinal occurrences in the Alps (Biella *et al*. 2017), and several other species naturally spreading into new areas (Bij de Vaate *et al*. 2002; Šima & Smetana 2012; Rutkowski *et al*. 2015). Although several species which are modifying their distributive ranges are supposed to be favoured or damaged by climate change, in most studies the association between climate and modification of distribution areas remains circumstantial and speculative (Nieto *et al*. 2014; Arbetman *et al*. 2017; Barbet-Massin *et al*. 2018; Pinkert *et al*. 2018). The results of our study suggest that the natural spreading of *B. haematurus* was favoured by warmer winters and specifically both warmer overwintering conditions (November-January) and also warmer months during the emergence of queens (February-March). A possible explanation of these results could be searched in a decreased mortality of queen in areas lying progressively northwards. That warmer winters are playing a role in the range expansion of species was already suggested for other species (Paradis *et al*. 2008; Caminade *et al*. 2012), but our study pointed out that a correlation exists between warmer winters (and not other seasons) and the northward natural spread of the studied species. Noteworthy, the current range expansion does not include some areas where warming of winters occurred. Possible reasons could to be searched in uneven sampling effort, absence of suitable corridors for colonization and of appropriate forested habitats. The latter factor could be detailed by means of an example; The Pannonian Plain is relatively close to the current *B. haematurus* range but most of it is not occupied by the species. Although warming winters happened in this area, the habitats there do not seem suitable for this bumblebee as that flat lowland region is predominantly used for intensive agriculture and only scattered forested remnants are preserved. As shown in our analysis, the climatic change is probably the most important trigger for expansion, but the interaction with components of species’ habitat requirements are still very relevant (Platts *et al*. 2019).

## Acknowledgements

The authors thank the institutions and the staff which contributed to accessing the data used in this study. We thank K. Horvath for the linguistic review. We warmly thank Irene Konovalova, Danilo Bevk, Paul Williams, Frederique Bakker, Maurizio Cornalba, Jakub Straka, and the participants at the ABIM – Alpine Bombus International Meeting for data sharing and/or for the fruitful discussions on *Bombus haematurus*.

## Contribution of authors

PB: project design and analysis; PB, AC, AG, JN, PS, MS, VS: data collection. All authors contributed to paper writing.

## Compliance with Ethical Standards

The authors declare that they have no conflict of interest and that no permission was required for the research.

## Supporting Information

Appendix S1 – A list of data providers used for building the dataset of the records for *Bombus haematurus*

